# Senataxin loss induces cGAS–STING-mediated mitochondrial dysfunction

**DOI:** 10.64898/2026.06.26.734838

**Authors:** Judith L.A. Fishburn, Hongchang Zhao, Will Fosselman, Julian Flores, Megan Wong, Tamira Singh, Steven Chen, Jacqueline Barlow

## Abstract

Ataxia with oculomotor apraxia type 2 (AOA2) is a rare neurodegenerative disease caused by loss-of-function mutations in *Senataxin,* which encodes an RNA:DNA helicase. Many studies on Senataxin loss focus on its putative roles in regulating transcription and RNA transcript localization. However, several phenotypes remain underexplored, including metabolic dysregulation associated with ataxias. Using Senataxin-deficient mouse cells, we observed increased nuclear and genomic instability, as well as innate immune activation via the cGAS-STING axis. We also observed elevated ROS levels, decreased mitochondrial function, and hyperfused mitochondria. Importantly, mitochondrial dysfunction depends on cGAS/STING activity, indicating that the two phenotypes are functionally connected. Senataxin-deficient mice and AOA2 patient cells similarly exhibit spontaneous innate immune activation, and AOA2 patient cells also show decreased mitochondrial function. Our work identifies a previously unseen phenotype for AOA2 in which loss of Senataxin results in mitochondrial dysfunction promoted by cGAS and STING.

**Summary:** This study demonstrates a previously unidentified phenotype in AOA2 and Senataxin research in which cGAS-STING promotes mitochondrial dysregulation in *Setx^-/-^* MEFs. Further, these innate immune activation and mitochondrial dysregulated phenotypes are recapitulated in AOA2 patient cells.

## Introduction

Ataxia with oculomotor apraxia type 2 (AOA2) is a recessive cerebellar ataxia that is estimated to affect 1 in 900,000 people globally (Anheim et al., 2010; Criscuolo et al., 2006) and shares clinical symptoms with other cerebellar ataxias, such as cerebellar degeneration, peripheral neuropathy, and loss of coordination (Anheim et al., 2009; Brusse et al., 2007). Disease onset for AOA2 patients typically occurs between 10 – 22 years of age, with rare cases diagnosed as early as 3. AOA2 patients also have a broad spectrum of symptoms that differ from those of other ataxias, including elevated serum alpha-fetoprotein (AFP) levels, slurred speech (Criscuolo et al., 2006), and sporadic oculomotor apraxia (Anheim et al., 2009; Criscuolo et al., 2006). There is no cure or clinically approved drug on the market to prevent or directly treat AOA2; patients manage symptoms through the use of a wheelchair for mobility, physical therapy, and educational tools such as computers to aid with speech and writing (Moreira & Koenig, 1993). AOA2 is caused by biallelic loss-of-function (LOF) mutations in the gene *Senataxin* (*SETX*) (Anheim et al., 2009; Brugger et al., 2014).

SETX is an RNA:DNA helicase that localizes primarily to the nucleus, with potential roles in regulating transcription (Miller et al., 2015; Suraweera et al., 2009; Yeo et al., 2015; Yüce & West, 2013), RNA splicing (Suraweera et al., 2009), antiviral response (Miller et al., 2015), meiosis (Yeo et al., 2015), and the DNA damage response (Hatchi et al., 2015; San Martin Alonso & Noordermeer, 2021; Yüce & West, 2013) presumably through its ability to resolve R-loops. R-loops are three-stranded nucleotide structures formed during active transcription when the nascent RNA re-anneals to its template strand, displacing the non-template strand as an ssDNA loop (Marnef & Legube, 2021; Thomas et al., 1976). Biochemical studies using the helicase domain of the yeast ortholog Sen1 or human SETX demonstrate their ability to resolve R-loops: the Sen1 helicase domain unwinds RNA:DNA hybrids with a preference for binding the DNA substrate (Han et al., 2017; Hasanova et al., 2023; Martin-Tumasz & Brow, 2015), while the helicase domain of human SETX resolves RNA:DNA hybrids with a preference for single-strand RNA (ssRNA) overhangs (Hasanova et al., 2023). Several studies indicate that depleting Senataxin by short hairpin or short interfering RNAs (shRNAs/siRNAs) increases R-loop levels (Cristini et al., 2019; Hatchi et al., 2015; Skourti-Stathaki et al., 2011; Yüce & West, 2013); however, other studies using knockout cells show no significant change in R-loop levels (Richard et al., 2020; Zhao et al., 2022). The direct role of Senataxin in transcription is also unclear due to conflicting studies examining transcription initiation (Kanagaraj et al., 2022), elongation (Han et al., 2025), and termination (Hasanova et al., 2023; Suraweera et al., 2009). Elevated DNA damage and accumulation of both single-strand and double-strand DNA breaks (ssDNA and dsDNA, respectively) have been identified in human cells lacking *SETX* (Becherel et al., 2015; Hatchi et al., 2015; Roda et al., 2014; Sollier et al., 2014). *SETX* deficiency is also associated with increased sensitivity to DNA-damaging agents, leading to oxidative stress and ssDNA breaks (Suraweera et al., 2007). Senataxin has been suggested to play a role in DNA repair by working with BRCA1 (Dutta et al., 2026; Hatchi et al., 2015); however, the specifics of this interaction, as well as its role in other DNA repair processes, remain incompletely understood. Thus, studies evaluating the direct role of Senataxin on R-loop homeostasis, transcription, and DNA repair are inconsistent, potentially due to the variability in how Senataxin is depleted or removed: acute depletion via degron (Han et al., 2025), siRNA/shRNA depletion (Crossley et al., 2023; Sberna et al., 2025), or knockout (Becherel et al., 2013; M. Li & Shao, 2024; Rao et al., 2024; Zhao et al., 2022). These concerns are exacerbated by methodological differences in measuring R-loops using DRIP, RDIP, SMRF-seq or R-ChIP (Chédin et al., 2021), and by the absence of a robust antibody against Senataxin.

Beyond R-loops, prior studies have investigated how the loss of Senataxin affects metabolic homeostasis, a shared feature of many inherited ataxias. Depletion of *SETX* expression leads to signs of dysregulated autophagy, mitochondrial function, and increased protein aggregates (Richard et al., 2020; Wen et al., 2024). A study of *Sen1* mutations showed decreased mitochondrial membrane potential and elevated ROS levels (Sariki et al., 2016). These elevated ROS levels have also been observed in human HeLa cells with *SETX* knockdown (Renaudin et al., 2021). Senataxin deficiency also activates the innate immune response (Crossley et al., 2023; Miller et al., 2015). Crossley et al., 2023 showed that *SETX-*depleted cells can induce innate immune activation via IRF3 activation. They suggested that initial activation was mediated by the RNA sensor TLR3 and the DNA sensor cGAS (Crossley et al., 2023). However, the contribution of these phenotypes to the etiology of AOA2 remains unclear.

To better understand the consequences of Senataxin loss associated with neurodegenerative phenotypes contributing to AOA2, including innate immune activation and metabolism, we investigated the impact of innate immune signaling on metabolic function. Using *Setx^-/-^* mouse embryonic fibroblasts (MEFs), we found elevated genomic and nuclear instability, activation of the cGAS-STING-IRF3 pathway, and increased interferon-stimulated gene (ISG) expression. We also observed increased ROS levels, reduced mitochondrial function, and differences in mitochondrial morphology, suggesting dysregulation of mitochondrial function and redox homeostasis. Interestingly, we found that cGAS and STING contribute to mitochondrial membrane potential defects but not morphological phenotypes. Finally, we observed increased innate immune activation in brain tissue from *Setx^-/-^* mice and in AOA2 patient fibroblasts. AOA2 patient cells also show mitochondrial dysfunction. Collectively, our work proposes a model where the loss of Senataxin in mammalian cells leads to mitochondrial dysfunction and innate immune activation, both driven by cGAS-STING.

## Methods

### Cell and mouse models

The *Setx^-/-^* mouse model was generated in a C57BL6/129Sv background and previously described in Becherel et al., 2013. From these mice, mouse embryonic fibroblasts (MEFs) were generated from day 13.5 embryos isolated from timed matings of *Setx^+/-^* mice, described in Lei, Y., 2013. Embryos were genotyped by PCR, as described in Becherel et al., 2013, and MEFs were immortalized by serial passaging. Human immortalized fibroblasts from a carrier and a patient with AOA2 were previously described in Fogel et al., 2014. All cells were cultured in fibroblast media (sterile-filtered DMEM, high glucose, no glutamine (Thermo Fisher Life Technologies), 10% fetal bovine serum, heat-inactivated (Thermo Fisher Life Technologies), and 1% penicillin-streptomycin (10,000 U/mL; Thermo Fisher Life Technologies). Cell lines were routinely tested (monthly to quarterly) for *Mycoplasma* and *Acholeplasma* species, common cell culture contaminants, using the MycoStrip 2.0 kit (InvivoGen).

### CRISPR knockout cell lines

Knockouts of *Cgas* and *Sting* in MEFs were generated by first selecting gRNAs using CRISPick (Doench et al., 2016; Sanson et al., 2018), followed by cloning gRNAs into the lentiCRISPRv2 puro plasmid (Addgene #52961) using a protocol established by the Zhang lab (Sanjana et al., 2014). The gRNAs used were *Cgas:* 5’-TGATAAGAAGTGTTACAGCA-3’, *Sting:* 5’-CAGTAGTCCAAGTTCGTGCG-3’, and Scramble (Scr): 5’-TATCTCGGAATCGAATGTTG-3’. The cloned gRNAs/lentiCRISPRv2 product was transfected into 293T cells using a calcium phosphate protocol with lentiviral packaging plasmids: 1.8 μg pMDLg/p-RRE, 1.25 μg pCMV-VSV-g, and 1.5 μg pRSV-Rev (J. H. Yu & Schaffer, 2006). After 48 hours, the supernatant was collected. Lentiviral particles were retrieved, and cell debris was removed by centrifugation and filtration through a 0.45 μm filter. The resulting lentiviral stocks were used to transduce WT and *Setx^-/-^* MEFs. Transduced cells were selected for puromycin resistance (4 μg/ml, ThermoFisher Scientific). *Cgas* and *Sting* knockouts underwent clonal selection via limiting dilution and were later verified by western blot; two clonal populations per knockout were selected and maintained.

### Real-time quantitative PCR (RT-qPCR)

Total RNA was extracted from MEFs, AOA2 patient-derived fibroblasts, and homogenized tissue using TRIzol (Invitrogen), followed by reverse transcription with the ProtoScript II First Strand cDNA Synthesis Kit (NEB) per the manufacturer’s protocol. qPCR was performed using 2X Universal SYBR Green Fast qPCR Mix (AB clonal) and analyzed on a Light Cycler 480 (Roche). The gene of interest was normalized to the housekeeping gene *Gapdh/GAPDH*, then to the WT controls using the standard ΔΔCT method. Mouse primers: *Gapdh* F: 5’-AATGTGTCCGTCGTGGATCT-3’ R: 5’- AGACAACCTGGTCCTCAGTG-3’, *Cxcl10* F: 5’-ATGACGGGCCAGTGAGAATG-3’ R: 5’-ATTCCGGATTCAGACATCTCT-3’, *Ccl5* F: 5’-ACGTCAAGGAGTATTTCTACAC-3’ R: 5’- GATGTATTCTTGAACCCACT-3’, *Ifit1* F: 5’-TCTAAACAGGGCCTTGCAG-3’ R: 5’- GCAGCCCTTTTTGATAATGT-3’, *Mfn1* F: 5’-ACAAGCTTGCTGTCATTGGG-3’ R: 5’-TCGACACTCAGGAAGCAGTT-3’, *Opa1* F: 5’-ATACTGGGATCTGCTGTTGG-3’ R: 5’-AAGTCAGGCACAATCCACTT-3’, *Fis1* F: 5’-AGATGGACTGGTAGGCATGG-3’ R: 5’- CTACAGGGGTGCAGGAGAAA-3’, and *Drp1* F: 5’-GTCGAGTCCCCATTCATTGC-3’ R: 5’-ACTACGACGATCTGAGGCAG-3’. Human primers: *GAPDH* F: 5’-GAAGGTGAAGGTCGGAGTCA-3’ R: 5’-AATGAAGGGCTCATTGATGG-3’, *CXCL1* F: 5’-GGAACAGAAGAGGAAAGAGAGAC-3’ R: 5’-TAGGACAGTGTGCAGGTAGA-3’, and *IFIT3* F: 5’-GCTGAGTCCTGATAACCAATACG-3’ R: 5’-GGCAAGGAGACTTTTCCAAGG-3’.

### Flow cytometry - TMRE

MEFs were grown overnight in a 6-well tissue-treated plate, and AOA2 cells were grown in 12-well tissue-treated plates (Genesee Scientific Corporation). Cells were treated with 1 μM oligomycin (Fisher Scientific) or 3 μM carbonyl cyanide-p-trifluoromethoxyphenylhydrazone (FCCP; Fisher Scientific) for 2 minutes, then treated with tetramethylrhodamine, ethyl ester (TMRE; Thermo Fisher Life Technologies) at 200 nM for MEFs and at 10 nM for human AOA2 and carrier fibroblasts for 30 minutes. Cells were washed with sterile-filtered 1X PBS, dissociated in 0.25% Trypsin-EDTA (Thermo Fisher Scientific), and neutralized in equal parts of the tissue culture media described above. Cells were transferred to Falcon 5 mL polystyrene round-bottom tubes (Fisher Scientific) and centrifuged at 1500 rpm for 5 minutes. Cells were then resuspended in 0.2% bovine serum albumin (BSA; ThermoFisher Scientific) in 1X PBS. Cells were then run on either a Canto II or an LSRII flow cytometer. Analysis was performed using FlowJo and gated on forward- and side-scatter area; from that gate, mean fluorescent intensity (MFI) was used as a measure of mitochondrial membrane potential.

### Immunofluorescence

Cells were seeded at 50-70% confluence and grown overnight on coverslips. Cells were washed three times with 1X PBS, fixed in 4% paraformaldehyde (PFA) in 1X PBS for 12 minutes at room temperature (RT), washed three times with 1X PBS, then permeabilized in 0.05% Triton X-100 in 1X PBS for 5 minutes at RT. Cells were washed three times with 1X PBS, then blocked in 2% BSA in 1X PBS for 1 hour. Cells were then incubated in primary antibodies for two hours, diluted in 1% BSA in 1X PBS at RT. Antibody dilutions were: mouse monoclonal anti-γ-H2AX (Millipore Sigma, clone JBW301) diluted at 1:1000, anti-lamin A/C (Cell Signaling Technologies, clone 4C11) at 1:150, anti-vinculin (Sigma Aldrich Inc., clone HVIN-1) at 1:1000, and anti-Tom20 (AB clonal, clone A19403) at 1:500. Cells were then washed three times with 1X PBS, then incubated 30 minutes to an hour in secondary antibodies diluted 1:100 to 1:500. Secondary antibodies used were anti-mouse Alexa Fluor 488 (ThermoFisher Life Technologies), anti-mouse Alexa Fluor 555 (ThermoFisher Life Technologies Corporation), and anti-rabbit Alexa Fluor 555 (ThermoFisher Life Technologies).

### Western blot

Cells were pelleted at 1500 rpm for 5 minutes, then washed in 1X PBS and pelleted again at 1500 rpm for 5 minutes. Cells were incubated in RIPA buffer (50 mM Tris–HCl, pH 8.0, 150 mM NaCl, 2 mM EDTA, pH 8.0, 1% NP-40, 0.5% sodium deoxycholate, 0.1% SDS, Protease Inhibitors) on ice for 30 minutes. Lysates were centrifuged at 1500 rpm for 5 minutes, and the supernatant was quantified for protein concentration using a Pierce BCA assay (Thermo Fisher Scientific). Whole-cell lysates (20 μg) were boiled in protein loading buffer for 10 minutes, placed on ice, loaded onto SDS-PAGE gels, and run under standard western blotting conditions. Transfer to a PVDF membrane was performed in standard transfer buffer on ice at a constant current of 330 mA for 1 hour, followed by blocking in 5% milk in 1X TBST for 1 hour. The primary antibodies used were anti-cGAS (Cell Signaling Technologies, clone D3O8O) at 1:1000, anti-STING (Cell Signaling Technologies, clone D1V5L) at 1:1000, anti-vinculin (Sigma Aldrich Inc., clone HVIN-1) at 1:2000, anti-OPA1 (Thermo Fisher Life Technologies Corporation, clone 1E8-1D9) at 1:1000, and anti-β-actin (AB clonal, clone AC026) at 1:10000. Secondary antibodies used were HRP-conjugated anti-mouse (AB clonal, clone AS003) and HRP-conjugated anti-rabbit (AB clonal, clone AS061) at 1:5000. Finally, Pierce ECL Western Blotting Substrate (Thermo Fisher Life Technologies Corporation) was used before imaging the membrane.

### Seahorse assays

The Seahorse XFe96 Bioanalyzer (Agilent Technologies) was used to perform the Mitochondrial Stress Test and Glycolytic Stress Test assays to measure oxygen consumption rate (OCR) and extracellular acidification rate (ECAR), as well as to extrapolate changes to oxidative phosphorylation and glycolytic rate (Amorim et al., 2022, 2024; Gillis et al., 2021). Briefly, cells were plated the night before, seeding 15,000 cells per well for AOA2 and carrier fibroblasts. For the mitochondrial stress test, 1 μM oligomycin, 3 μM FCCP, and 0.5 μM each of rotenone and antimycin A (Fisher Scientific Company) were used. For the glycolytic stress test, 25 mM glucose (Agilent Technologies), 1 μM of oligomycin, and 50 mM 2-deoxy-D-glucose (2-DG; Fisher Scientific Company) were used.

### pH-Xtra Glycolysis Assay

The pH-Xtra Glycolysis Assay (Agilent Technologies) was used to measure ECAR rates. In a 96-well tissue-treated plate (Genesee Scientific Corporation), MEFs were plated overnight and seeded at a density of 10,000 cells/mL. The assay was performed according to Agilent’s pH-Xtra Glycolysis Assay. Briefly, cells and assay material were incubated at 37°C in a non-CO_2_ incubator with a water bath for at least 30 minutes. Controls used were wells without the pH-Xtra reagent or cells; wells with only the pH-Xtra reagent; a cell-free positive control with the pH-Xtra reagent and 100 μg/mL Glucose Oxidase (Fisher Scientific Company); and a cell-free negative control with 50 mM 2-DG. Cells in experimental wells were treated with the pH-Xtra reagent alone, pH-Xtra reagent plus 100 μg/mL Glucose Oxidase (Fisher Scientific Company), pH-Xtra reagent plus1 μM oligomycin (Fisher Scientific Company), or pH-Xtra reagent plus 3 μM FCCP (Fisher Scientific Company). Assays were run on the SpectraMax iD5 (Molecular Devices) with the plate read every 30 seconds at an excitation of 350 ± 60 nm and an emission of 616 ± 10 nm for 2 hours. ECAR rates were normalized to WT, and the fold change was plotted.

### Dual Luciferase Reporter Assay

The dual-luciferase reporter assay for IRF3 and NF-κB activity was performed using the Promega Dual-Luciferase Reporter Assay System (E1910) and protocol. Cells were seeded into 24-well plates and cultured until they reached approximately 70–80% confluency. The cells were then co-transfected with the indicated firefly luciferase reporter plasmids together with the TK-Renilla luciferase plasmid as an internal transfection control (DeFilippis et al., 2010; H. Yu et al., 2022, 2025). At 48 hours post-transfection, cells were washed with 1X PBS and lysed using passive lysis buffer. The cell lysates were collected, and firefly and Renilla luciferase activities were measured using a GloMax luminometer (Promega) with a microinjector. Firefly luciferase activity was normalized to Renilla luciferase activity to correct for transfection efficiency. The relative promoter activity was calculated and compared with that of the control plasmid-transfected group. The results were expressed as relative luciferase activity.

### HRP-Luminol ROS Assay

The HRP-luminol assay was adapted from Hong, Z. et al. 2016 and was performed on WT and *Setx^-/-^* MEFs (Zhu et al., 2016). Cells were plated at 100,000 cells per well, then centrifuged in a black well of a clear-bottom plate (Corning). Cells were treated with 10 μg/mL peroxidase from horseradish (HRP; Millipore Sigma), 10 μM luminol sodium salt (Millipore Sigma-Aldrich), 20 μM ß-Lapachone (Millipore Sigma), 100,000 U/mL catalase from bovine liver (Millipore Sigma-Aldrich), and 25 mM sodium pyruvate (ThermoFisher Life Technologies). Then the plate was run on a GloMax luminometer (Promega).

## Results

### *Setx^-/-^* MEFs exhibit increased genome instability and nuclear abnormalities

To evaluate genome instability in Senataxin-deficient cells, we first quantified micronucleus (MN) formation in immortalized *Setx^-/-^* MEFs derived from mice harboring a germline deletion of Senataxin (Becherel et al., 2013). We observed a significantly higher fraction of cells with MN in *Setx^-/-^* MEFs than in WT controls (Fig. 1, A and B). MN formation can be caused by DNA damage or mitotic errors, leading to lagging chromosomes or chromosome fragments that are not incorporated into the main nucleus as the nuclear envelope (NE) is re-established in anaphase (Krupina et al., 2021). To determine if the main nucleus also exhibited genome instability, we measured γ-H2AX foci, the phosphorylated form of histone H2AX that accumulates at ssDNA and dsDNA breaks (Merighi et al., 2021). We found that *Setx^-/-^* MEFs have a higher percentage of cells with 10 or more γ-H2AX foci than WT cells, indicating significantly more DNA damage (Fig. 1, A and C). This data suggests that increased DNA damage may promote MN formation and genome instability in *Setx^-/-^* MEFs.

**Figure 1:**
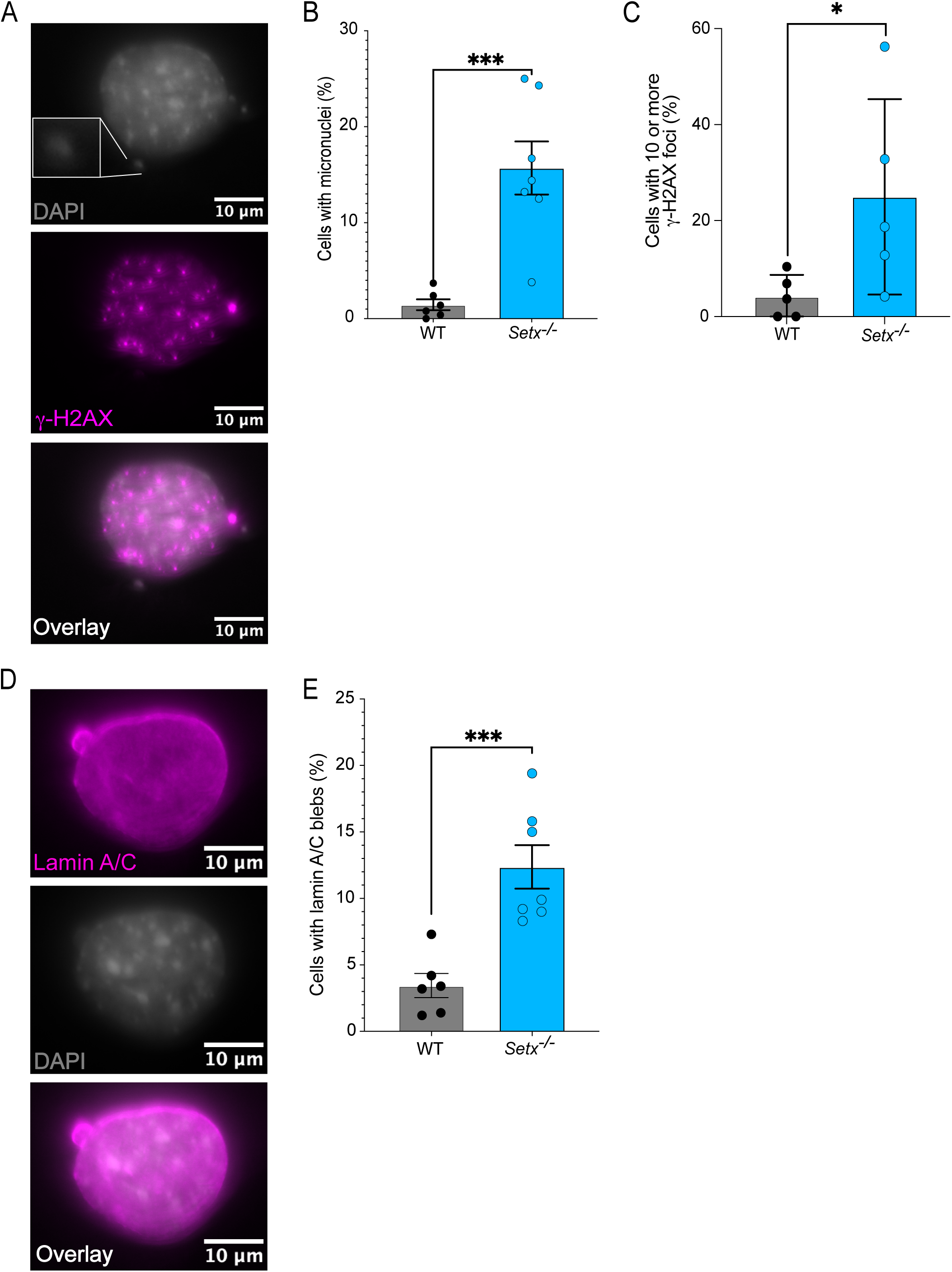
Elevated micronuclei, γ-H2AX foci, and lamin A/C blebbing events Setx-/- MEFs. (A) Representative image of a *Setx^-/-^* cell stained for DNA damage with DAPI in greyscale and γ-H2AX in magenta; a pop-out box highlights a micronucleus (MN) in the top panel and magenta-bright spots indicative of γ-H2AX foci in the bottom two panels. **(B)** Percent of cells with MN in WT and *Setx^-/-^* MEFs (n=7 biological replicates). The error bars show the standard error of the mean (SEM). **(C)** The percent of cells with 10 or more γ-H2AX foci (n=7 biological replicates). **(D)** Representative images of DAPI (grey) and lamin A/C (magenta) in a *Setx^-/-^* cell. The top and overlay panels show lamin A/C blebbing, which appears as a bubble coming off the nucleus. **(E)** The percentage of cells with lamin A/C blebs in WT and *Setx^-/-^* MEFs (n=7 biological replicates). The confidence intervals (CI) for each figure with statistics were set to 95%, and error bars represent the SEM. All statistics were analyzed using two-tailed unpaired t-tests. P values: * = p < 0.05; *** = p < 0.001.

Nuclear instability due to dysfunctional nuclear structural proteins can also contribute to MN formation (Schultz et al., 2025). Lamin proteins play critical roles in the structure and stability of the nucleus by forming a meshwork beneath the inner nuclear membrane of the NE, influencing chromosome organization, cell motility, gene expression, and regulation (Dubik & Mai, 2020; Schultz et al., 2025). Lamin defects are associated with nuclear instability, DNA damage accumulation, and nuclear ruptures that can contribute to MN formation; mutations in lamin genes can lead to diseases like progeria and cancer (Chojnowski et al., 2015; Chu et al., 2025; Duan et al., 2026; Dubik & Mai, 2020, 2020; Zwerger et al., 2013). To determine whether the NE and lamin structure are altered, we measured the percentage of cells with lamin A/C blebbing events. Blebbing occurs when the NE becomes unstable and forms bubble-like structures (Fig. 1 D). We found that the percentage of cells exhibiting lamin A/C blebbing was significantly higher in *Setx^-/-^*MEFs (Fig. 1 E). Thus, loss of Senataxin leads to an increase in *γ*-H2AX foci, lamin A/C blebbing, and MN, demonstrating genomic and nuclear instability.

### Innate immune activation occurs through the cGAS-STING axis in *Setx^-/-^* MEFs

Genomic and nuclear instability can trigger innate immune activation, thereby initiating the inflammatory response. One way this occurs is activation of the cytosolic dsDNA sensor, cGAS (Mackenzie et al., 2017). Work from Crossley et al. suggests that the innate immune system is activated in *SETX*-depleted HeLa cells via TLR3 and cGAS-STING (Crossley et al., 2023). To determine whether the cGAS/STING pathway is spontaneously induced in *Setx*-knockout MEFs, we examined total levels of cGAS and STING proteins. We found that both cGAS and STING protein levels were increased in *Setx^-/-^* MEFs (Fig. 2 A). Increased expression of cGAS and STING is associated with Alzheimer’s disease (Hou et al., 2021). Further, their activity is implicated in ataxia telangiectasia (A-T), a cerebellar ataxia caused by mutations in the DNA damage and checkpoint kinase ATM (Aguado et al., 2021).

**Figure 2:**
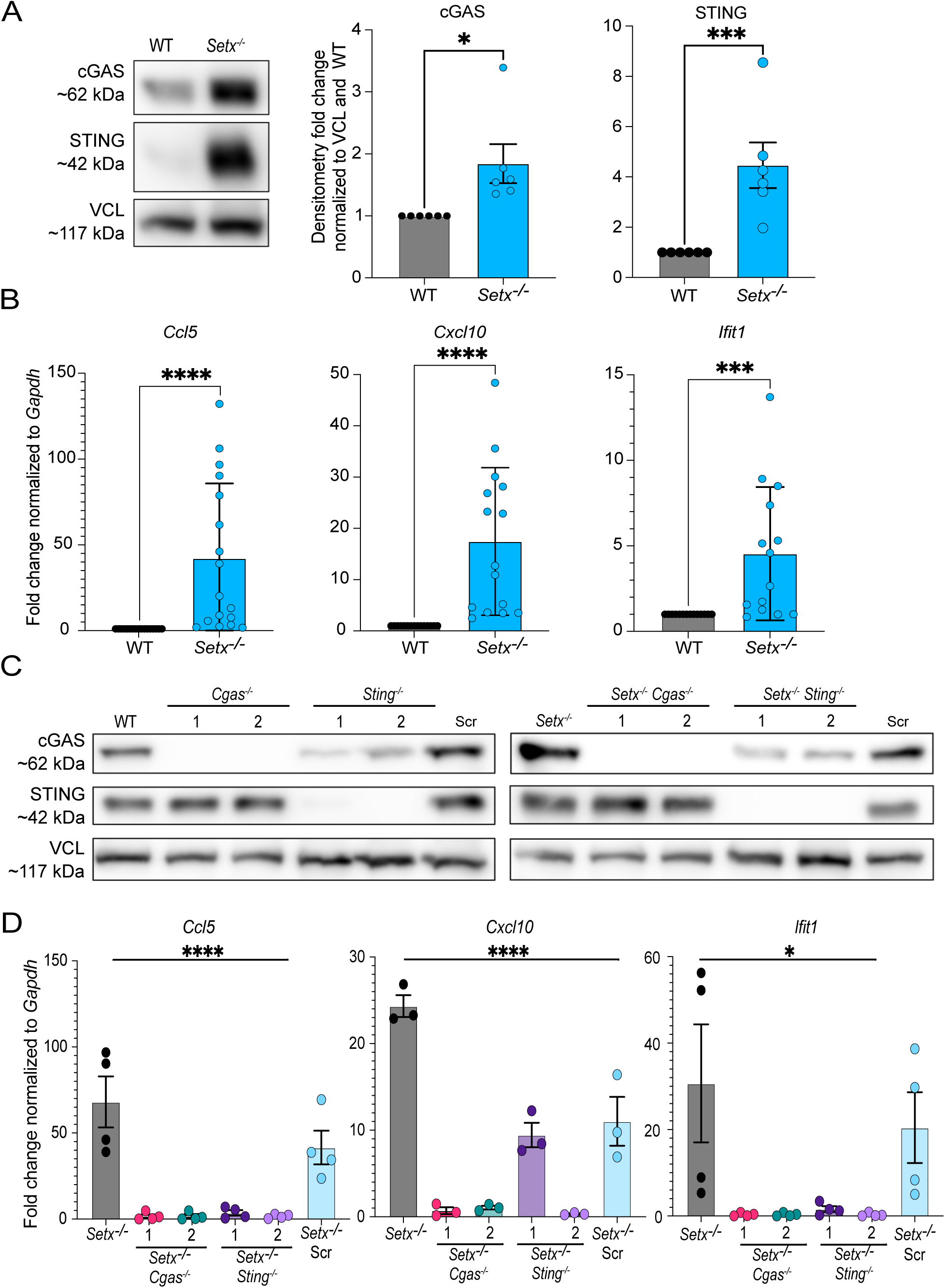
Innate immune activation in *Setx^-/-^* MEFs through the cGAS-STING axis. **(A)** Representative western blot of WT and *Setx^-/-^* MEF protein lysates with antibodies against cGAS (∼62 kDa), Sting (∼42 kDa), and Vinculin (VCL) (∼117 kDa). Densitometry for cGAS and STING are in the right two panels (n=6). **(B)** Interferon-stimulated gene (ISG) expression of *Ccl5, Cxcl10,* and *Ifit1* measured via RT-qPCR (n=15-17). **(C)** Representative western blots of WT, *Cgas^-/-^*, *Sting^-/-^*, Scramble (Scr), *Setx^-/-^*, *Setx^-/-^Cgas^-/-^*, *Setx^-/-^Sting^-/-^*, and *Setx^-/-^* Scr MEFs probed with antibodies to cGAS (∼62 kDa), Sting (∼42 kDa), and Vinculin (VCL) (∼117 kDa). Two independent single-clone populations for each of *Cgas^-/-^* and *Sting^-/-^* were generated and labeled 1 or 2. **(D)** ISG expression of *Ccl5, Cxcl10,* and *Ifit1* measured via RT-qPCR in all MEF lines shown in (C) (n=4). The CI was set to 95%, and error bars represent the SEM. Two-tailed unpaired t-tests (A and B) and One-way ANOVA (D); P values: * = p < 0.05; *** = p < 0.001; ****= p <0.0001.

Exogenous DNA-damaging agents or loss of DNA repair factors predominantly induce a type I IFN response (Brzostek-Racine et al., 2011; Härtlova et al., 2015; Sui et al., 2017; Taffoni et al., 2021). To determine whether the type I IFN signaling cascade is induced, we next measured interferon-stimulated gene (ISG) expression and quantified mRNA transcript levels of the cytokines *Ccl5* and *Cxcl10* and the antiviral factor *Ifit1* by RT-qPCR. We found that *Ccl5, Cxcl10,* and *Ifit1* expression were all significantly increased in *Setx^-/-^* MEFs (Fig. 2 B), indicating activation of the type I IFN response. We next evaluated if cGAS and STING are required for ISG expression by generating two independent single-clone knockouts for *Cgas* and *Sting* in WT and *Setx^-/-^* MEFs using CRISPR-Cas9 (Fig. 2 C). We found that knockout of *Cgas* or *Sting* significantly reduced ISG expression of *Ccl5, Cxcl10*, and *Ifit1* in *Setx^-/-^* MEFs but had no significant impact on ISG expression in WT cells (Fig. 2 D, Fig. S1 A). These results suggest that both cGAS and STING are essential for inducing the type I IFN response seen in *Setx^-/-^* MEFs.

To determine which transcription factors downstream of cGAS-STING are active, we used a dual luciferase assay to detect IRF3 (DeFilippis et al., 2010) and NF-κB activity (H. Yu et al., 2022, 2025). We observed a significant increase in IRF3 activity, whereas NF-κB activity was modestly but not significantly increased (Fig. S1 B). These results indicate that IRF3 is the predominant transcription factor driving ISG expression. Combined, this work supports previous findings from Crossley et al. that cGAS-STING-IRF3 signaling induces innate immune activation in Senataxin*-*deficient cells.

### *Setx^-/-^* MEFs have increased ROS and altered metabolism

Induction of the type I IFN response and STING activation can induce reactive oxygen species (ROS) production and accumulation through several mechanisms (Hayman et al., 2021; Kumova & Carey, 2020). ROS are important in type I IFN signaling (Hwang et al., 2021; Wu et al., 2024), are thought to play a role in the etiology of Friedreich’s ataxia (FA) (Lupoli et al., 2018; Schulz et al., 2000), and are a source of oxidative stress, a hallmark of neurodegeneration (Wilson et al., 2023). Therefore, we investigated whether *Setx^-/-^* MEFs exhibit increased ROS levels. Indeed, we found that *Setx^-/-^*MEFs exhibit increased ROS levels as measured by a luminol-horseradish peroxidase chemiluminescence assay (Zhu et al., 2016), which primarily detects hydrogen peroxide (H_2_O_2_; Fig. 3 A). The increased ROS can be reduced by adding the H_2_O_2_ scavenger catalase. ROS can increase with H_2_O_2_ treatment, further indicating that the increase in ROS measured is likely due to H_2_O_2_ (Fig. S2).

**Figure 3:**
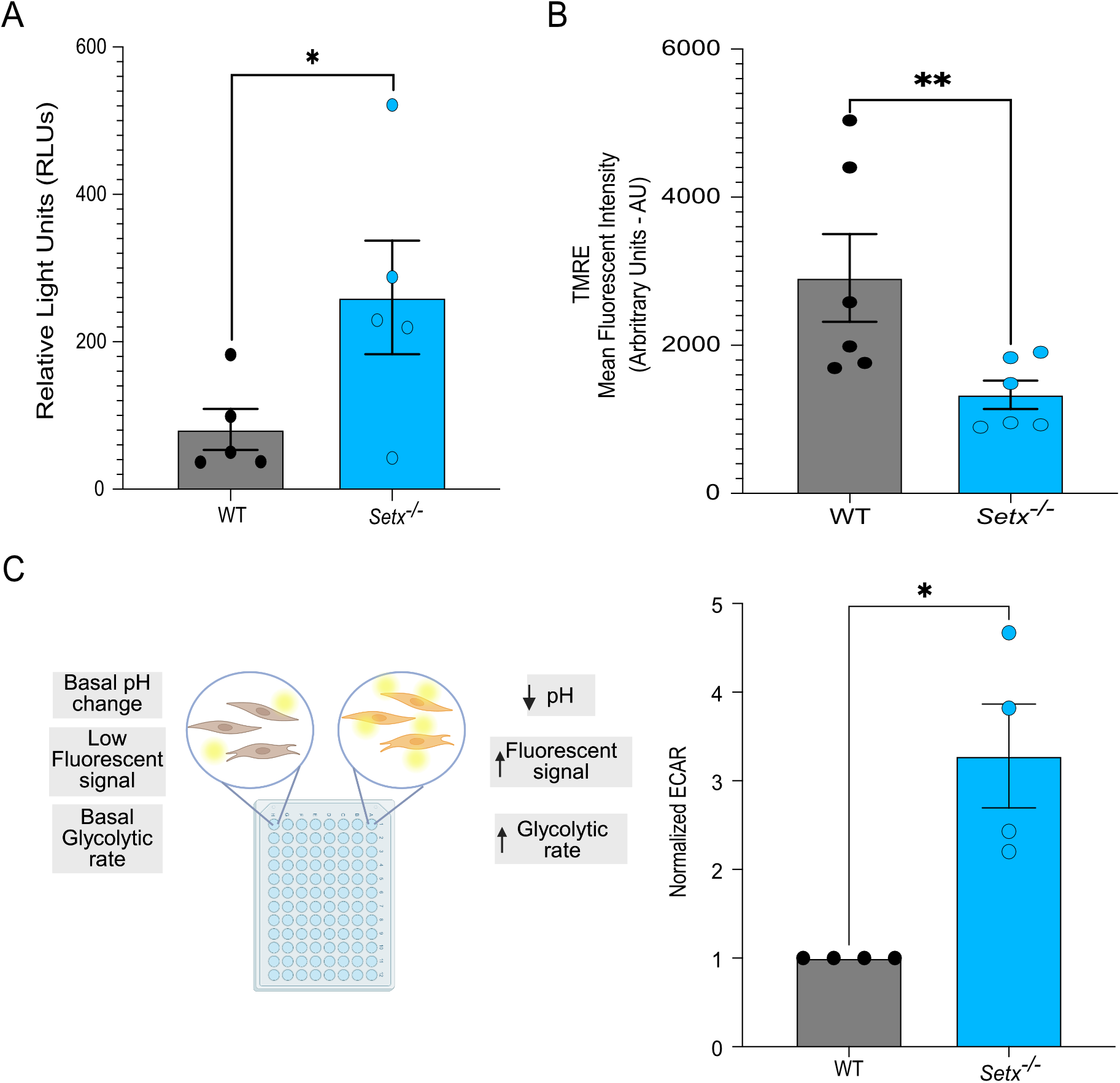
Mitochondrial membrane potential, ROS, and ECAR are altered in Setx-/ MEFs. **(A)** ROS levels measured in WT and *Setx^-/-^* MEFs where luminescent signal in an HRP-luminol assay increases as ROS levels do (n=5). **(B)** Quantitation of TMRE signal in *Setx^-/-^* and WT MEFs (n=6). **(C)** Left panel: Diagram of the pH-Xtra assay, which uses a proprietary dye from Agilent Technologies that emits light at ∼616 nm when pH becomes more acidic. Right panel: Extracellular acidification rate (ECAR) was normalized to WT (n=4). The CI was set to 95%, and error bars represent the SEM. (A and B) Two-tailed unpaired t-tests. P values are as follows: * = p < 0.05; ** = p < 0.01; ****= p <0.0001.

Activation of cGAS-STING and increased ROS levels also influence mitochondrial function (Kim et al., 2023; Zhang et al., 2025). Studies of A-T, caused by mutations in the DNA repair protein kinase ATM, show that both mouse- and patient-derived organoid models exhibit elevated oxidative stress and mitochondrial dysregulation (Leeson et al., 2024; Valentin-Vega et al., 2012). To determine whether loss of Senataxin similarly causes mitochondrial defects, we measured changes in mitochondrial membrane potential using the TMRE assay. We found that *Setx^-/-^* MEFs have significantly lower TMRE signal than WT cells (Fig. 3 B), indicating reduced mitochondrial membrane potential and mitochondrial dysregulation. In response to mitochondrial dysregulation, MEFs can compensate for energy demands by increasing glycolytic activity (H. Li et al., 2019). Further, models of spinocerebellar ataxia type 1 (SCA1) exhibit mitochondrial dysregulation and increased extracellular acidification rate (ECAR), an indicator of glycolytic rate (Ford-Roshon et al., 2025). We measured ECAR and found that *Setx^-/-^* MEFs also exhibit increased ECAR compared to WT cells (Fig. 3 C). Based on these results, we propose that Senataxin deficiency increases oxidative stress and dysregulates mitochondrial function, causing cells to shift their metabolism toward glycolysis.

### *Setx^-/-^* MEFs display hyperfused mitochondria

Mitochondrial dysfunction is frequently coupled with morphological changes; in primary mouse hepatocytes treated with acetaminophen, mitochondrial membrane potential was reduced and mitochondrial fragmentation increased (Umbaugh et al., 2021). Endogenous stress, such as from oxidative signaling, can also induce mitochondrial fragmentation (Fu et al., 2020). However, stress-induced mitochondrial hyperfusion (SIMH) can also occur in response to stressors such as UV irradiation, actinomycin D, serum deprivation, or amino acid deprivation (Tondera et al., 2009). We next evaluated differences in mitochondrial morphology using confocal microscopy of the outer mitochondrial membrane protein Tom20 and the membrane-cytoskeletal protein Vinculin to visualize the cell membrane. We found that *Setx^-/-^* MEFs have distinctly elongated mitochondrial morphology from WT MEFs, suggestive of more fused mitochondria (Fig. 4 A). Based on previous reports, we classified mitochondrial morphology into two groups: either punctate (fragmented) or intermediate and elongated/filamentous mitochondria (Giedt et al., 2016; Picard et al., 2013). A significant proportion of mitochondria in WT cells were punctate or intermediate (Fig. 4 B), consistent with previous studies (De Gaetano et al., 2020; R. Wang et al., 2021). However, a significantly larger fraction of *Setx^-/-^* cells exhibited a hyperfused and filamentous mitochondrial morphology (Fig. 4 C). *Setx^-/-^*MEFs were not significantly different in both cellular and mitochondrial volume compared to WT MEFs; the ratio of mitochondrial to cellular volume was similar between *Setx^-/-^* and WT MEFs (Fig. S3, A and B). The mean fluorescent intensity of TOM20 was also similar between WT and *Setx^-/-^*(Fig. S3 C). Taken together with our observations that *Setx^-/-^*MEFs exhibit increased inflammation, elevated ROS, and metabolic dysregulation, the altered mitochondrial morphology suggests that Senataxin loss provokes a SIMH state.

**Figure 4:**
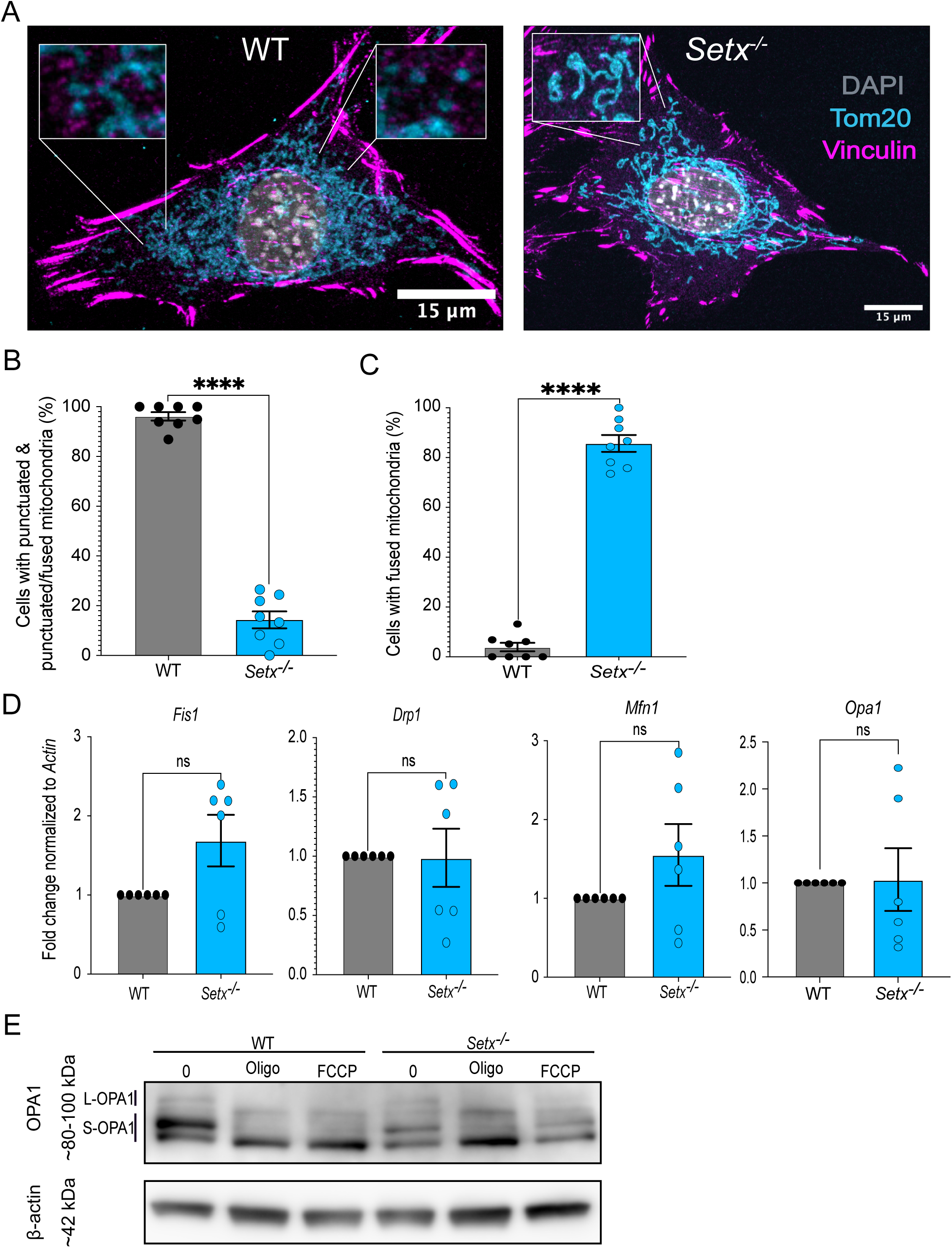
Mitochondrial hyperfusion in *Setx^-/-^* MEFs. **(A)** Representative images of WT and *Setx^-/-^* MEFs labeled with DAPI (nuclear DNA, grey), Tom20 (mitochondria; cyan), and Vinculin (cell membrane; magenta). **(B)** Quantification of punctated and punctated/fused (intermediate) mitochondria in WT and *Setx^-/-^* MEFs (n=8). **(C)** Quantitation of fused mitochondria in WT and *Setx^-/-^* MEFs (n=8). **(D)** *Fis1* and *Drp1* (fission genes), and *Mfn1* and *Opa1* (fusion genes) gene expression using RT-qPCR in WT and *Setx^-/-^*MEFs (n=6). **(E)** Western blot of WT and *Setx^-/-^*MEFs with or without 1 μM oligomycin (oligo) or 3 μM FCCP. OPA1 isoforms are shown with vertical lines indicating where the 2 L-OPA1 isoforms and the 3 S-OPA1 isoforms are. Two-tailed unpaired t-tests with a 95% CI were used. P values: **** = p < 0.0001; ns = not significant.

Multiple proteins can drive morphological changes in mitochondria between fusion and fission. Mitofusin 1 and 2 (MFN1 and MFN2) and OPA1 regulate fusion of the outer and inner membranes of mitochondria, respectively (Cipolat et al., 2004; Gao & Hu, 2021). Key regulators of fission include DRP1, which binds to the outer membrane of mitochondria, and FIS1, an adaptor protein that facilitates fission (Fonseca et al., 2019; Zerihun et al., 2023). To determine if these regulators are altered in *Setx^-/-^*MEFs, we first measured changes in *Mfn1, Opa1, Drp1,* and *Fis1* gene expression and found no significant difference (Fig. 4 D). Posttranslational modifications also influence mitochondrial morphological differences, specifically OPA1 cleavage (R. Wang et al., 2021). OPA1 has several protein isoforms arising from alternative splicing and posttranslational cleavage by the OMA1 and YME1L proteases (R. Wang et al., 2021). The two unprocessed OPA1 isoforms are referred to as long-OPA1 (L-OPA1), and the three cleaved isoforms are referred to as short-OPA1 (S-OPA1) (Gilkerson et al., 2021). The prevalence of isoforms can vary under stress, thereby influencing mitochondrial morphology (Gilkerson et al., 2021; X. Li et al., 2022; Mishra et al., 2014; R. Wang et al., 2021). We measured the abundance of L-OPA1 and S-OPA1 isoforms in WT and *Setx^-/-^* MEFs by western blot. The ATP synthase inhibitor oligomycin (oligo) and carbonyl cyanide-p-trifluoromethoxy-phenylhydrazone (FCCP), an ionophore that uncouples the mitochondrial membrane potential, both induce L-OPA1 cleavage and were used as positive controls for S-OPA1 isoform enrichment (MacVicar & Lane, 2014; X. Wang et al., 2014). Consistent with prior reports, treatment with either FCCP or oligo increases the abundance of S-OPA1 fragments (Fig. 4 E). We found that *Setx^-/-^* MEFs have less OPA1 expression, similar levels of S-OPA1 isoforms, and reduced levels of some of the L-OPA1 isoforms (L-OPA1 labeled on blots; Fig. 4 E). Taken together, this data suggests that OPA1 cleavage and altered isoform abundance may be driving the mitochondrial morphology differences observed in *Setx^-/-^* MEFs.

### Knockout of Cgas or Sting increases mitochondrial membrane potential but does not influence mitochondrial morphology in *Setx^-/-^* MEFs

We next examined whether cGAS and STING contribute to the increased DNA damage, nuclear instability, or altered mitochondrial morphology observed in *Setx^-/-^* MEFs. Here, we found that loss of either cGAS or STING had no impact on WT TMRE signal but increased TMRE in *Setx^-/-^* cells, with 3 out of 4 clones showing a significant increase (Fig. 5, A and B). These results indicate that cGAS/STING signaling reduces mitochondrial membrane potential in *Setx^-/-^* MEFs. However, we found that loss of cGAS or STING had no significant impact on mitochondrial morphology in either WT or *Setx^-/-^* MEFs (Fig. 5 C). We observed a reduction in MN in *Setx^-/-^Cgas^-/-^* cells, but the reduction was significant only for one clone (Fig. 5 D). The amount of nuclear blebbing was not significantly reduced in *Setx^-/-^* MEFs upon cGAS/SITNG loss (Fig. 5 E). These results show that deletion of cGAS or STING does not reverse mitochondrial morphology or genome instability phenotypes in MEFs, suggesting that these phenotypes may be separate phenomena.

**Figure 5:**
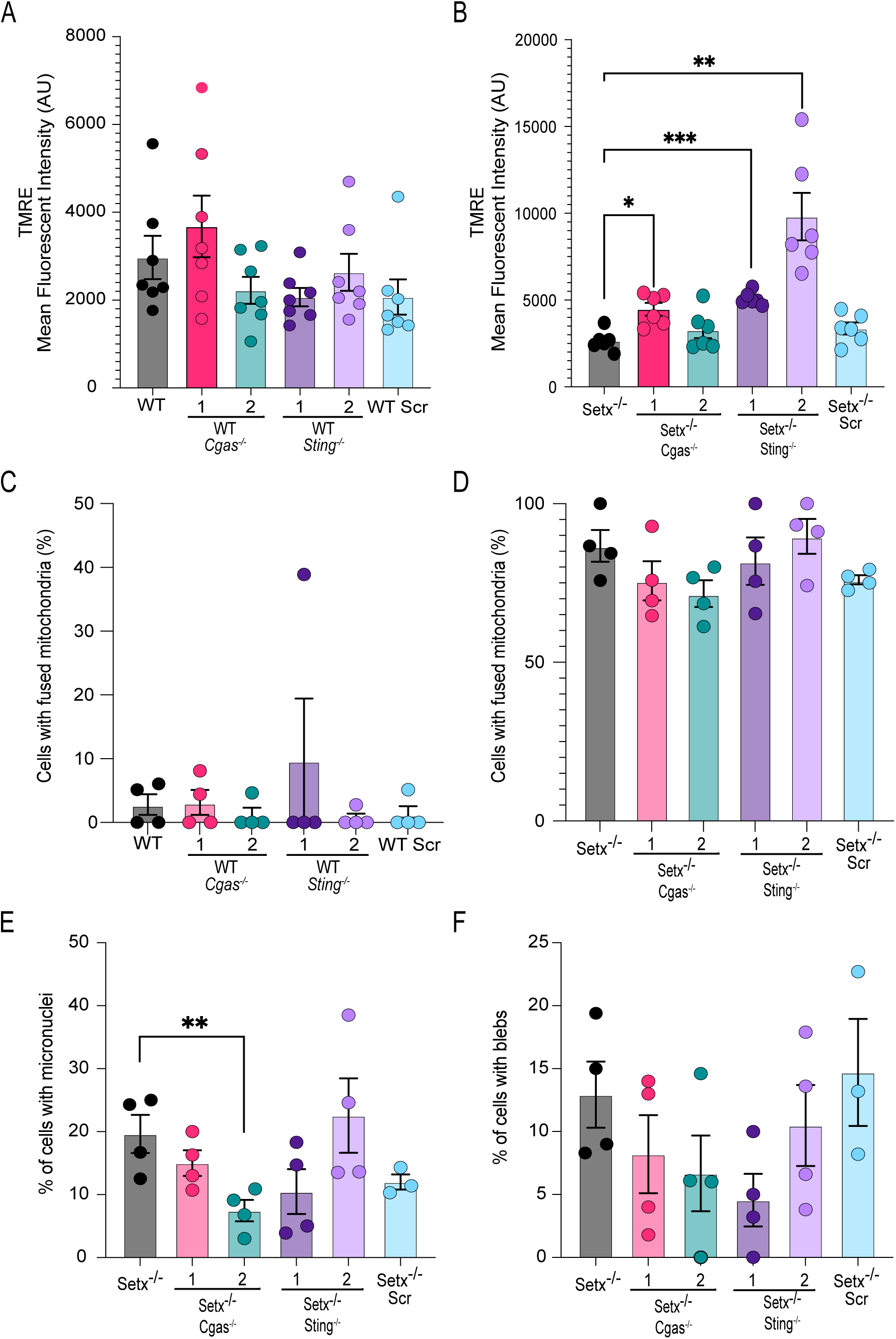
The mitochondrial membrane potential in *Setx^-/-^ Cgas^-/-^* and *Setx^-/-^ Sting^-/-^* MEFs. **(A)** *Cgas^-/-^*, *Sting^-/-^*, and Scr MEFs: mitochondrial membrane potential evaluated using TMRE and quantified by MFI (n=7). **(B)** *Setx^-/-^ Cgas^-/-^, Setx^-/-^ Sting^-/-^*, and *Setx^-/-^*Scr MEFs TMRE MFI representing mitochondrial membrane potential signal (n=6). **(C)** Quantitation of mitochondrial morphology in WT, *Cgas^-/-^* and *Sting^-/-^* MEFs (n=4). **(D)** Quantitation of mitochondrial morphology in *Setx^-/-^Cgas^-/-^, Setx^-/-^Sting^-/-^* and *Setx^-/-^* MEFs (n=4). **(E)** The percentage of *Setx^-/-^ Cgas^-/-^* and *Setx^-/-^ Sting^-/-^* cells with micronuclei (n=4). **(F)** The percentage of cells with lamin A/C blebs in *Setx^-/-^ Cgas^-/-^* and *Setx^-/-^ Sting^-/-^*cells (n=4). Two independent single-clone populations for each of *Cgas^-/-^* and *Sting^-/-^* labeled 1 or 2. One-way ANOVA. P values are as follows: * = p < 0.05; ** = p < 0.01; *** = p < 0.001.

### ISG expression is elevated in the brains of *Setx^-/-^* mice and AOA2 patient cells

Given the increase in ISG expression in MEFs, we next evaluated ISG expression in primary tissues from WT and *Setx^-/-^* mice. We found that the *Setx^-/-^* mice have higher ISG expression in the brain, with *Cxcl10* expression being significantly increased (Fig. 6 A). This increase appears tissue-specific, as other tissues, such as the heart and kidney, do not show a significant increase in ISG expression (Fig. S4, A and B). These results suggest that loss of Senataxin increases innate immune gene expression associated with inflammation in the brains of mice.

**Figure 6:**
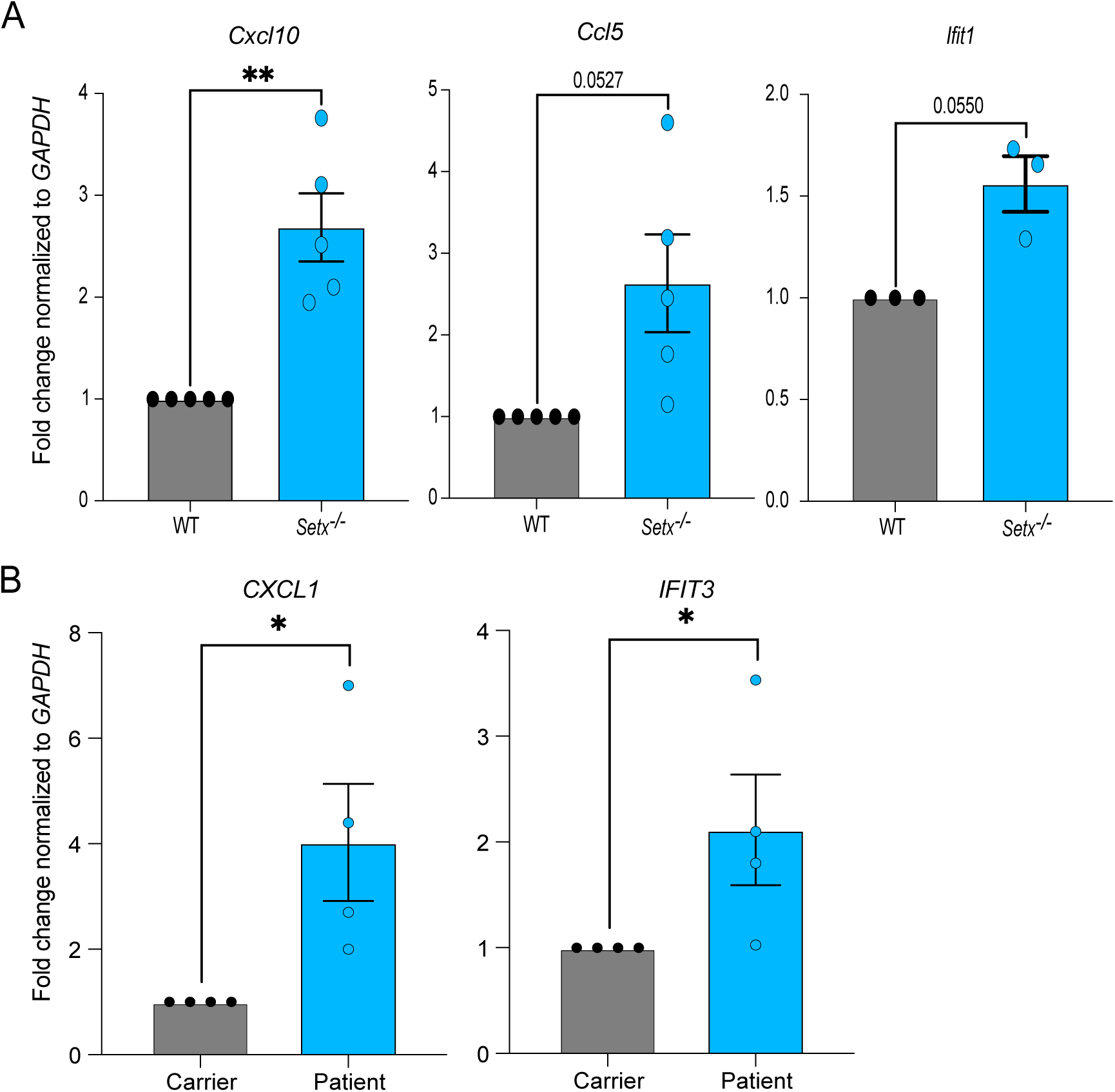
ISG expression in *Setx^-/-^* mouse brains and AOA2 patient-derived fibroblasts. **(A)** RNA isolated from whole brains of both WT and *Setx^-/-^* mice and measured for changes in *Cxcl10*, *Ccl5,* and *Ifit1* expression using RT-qPCR (n=3-5). **(B)** ISG expression of *CXCL1* and *IFIT3* measured using RT-qPCR in AOA2 patient-derived fibroblasts (n=4). Two-tailed unpaired t-tests with a 95% CI. P values are as follows: * = p < 0.05; ** = p < 0.01.

To determine if ISG expression is also altered in human cells lacking Senataxin, we next measured ISG expression in AOA2 patient-derived fibroblasts. Using immortalized fibroblasts from a carrier and an AOA2 patient (Fogel, Cho, et al., 2014), we measured ISG expression of two type I IFN genes, *CXCL1* and *IFIT3*. CXCL1 is a chemokine, and IFIT3 plays a role in the antiviral response and enhances overall innate immune activation (Glasner et al., 2025; Tang et al., 2026). We found that *CXCL1* and *IFIT3* expression are both increased in AOA2 compared to carrier fibroblasts (Fig. 6 B), consistent with MEFs and primary mouse brain tissue. Together, these results indicate that the absence of Senataxin in both mouse and human cells induces an innate immune response that typically leads to inflammation. Further, the brain-specific increase of ISG expression in mice may partially explain the tissue-specific impacts of germline SETX mutations.

### AOA2 patient cells have decreased mitochondrial function

We next examined whether patient AOA2 cells exhibited mitochondrial dysregulation like *Setx^-/-^*MEFs. Indeed, we found that AOA2 fibroblasts exhibit a significantly lower TMRE signal than carrier cells, indicating reduced mitochondrial membrane potential (Fig. 7 A). Of note, carrier cells have one inactive *SETX* allele and have been shown to have reduced SETX protein levels (Fogel, Lee, et al., 2014), which may explain why the reduction in TMRE is less pronounced than that observed in MEFs (Fig. 3 B). Using the Seahorse metabolic analyzer, we found that patient AOA2 cells also show lower mitochondrial respiratory capacity (Fig. 7, B – D; Fig. S5, A and B). AOA2 patient cells also tended to have higher basal ECAR and glycolytic reserve, but these differences were not significant (Fig. 7, E-G). Glycolysis and glycolytic capacity were similar between carrier and patient cells (Fig. S5, C and D). Together, these results indicate that AOA2 patient-derived cells exhibit defective mitochondrial function, like that of *Setx^-/-^* MEFs, and that this mitochondrial dysfunction likely alters other metabolic pathways.

**Figure 7:**
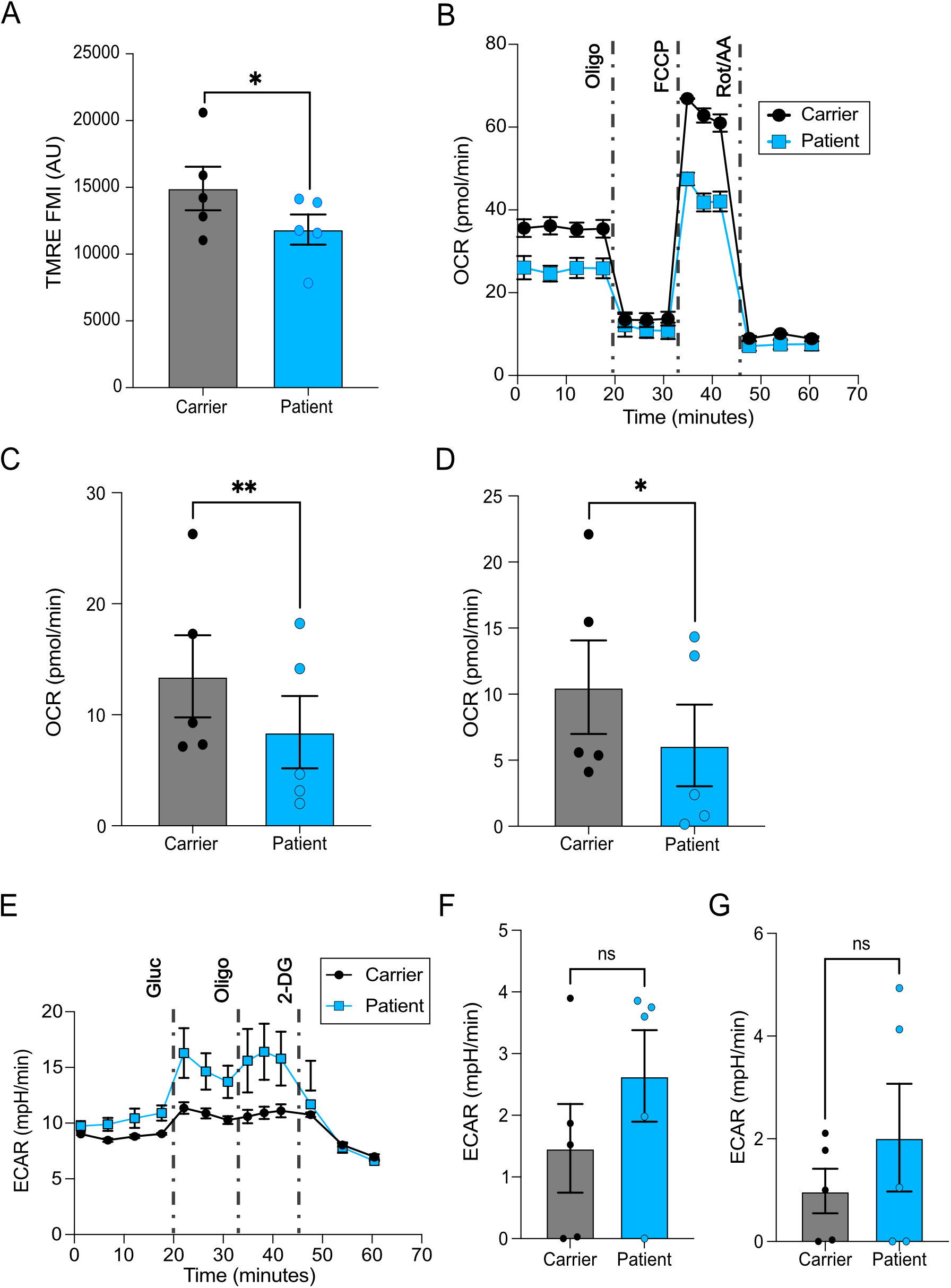
Mitochondrial function is dysregulated in AOA2 patient cells. **(A)** Quantitation of mitochondrial membrane potential by TMRE in AOA2 patient-derived fibroblasts and carrier fibroblasts (n=5). **(B)** Image of an optimal Seahorse Mito Stress Test assay comparing AOA2 patient cells to carrier cells, where the y-axis is oxygen consumption rate (OCR) and the x-axis is time in minutes. Note the drugs used at different time points: oligomycin (oligo), carbonyl cyanide-p-trifluoromethoxyphenylhydrazone (FCCP), and rotenone plus antimycin A (Rot/AA). (**C)** Quantitation of the basal oxygen consumption rate (OCR) from Mito Stress Test assay in AOA2 patient and carrier cells (n=5). **(D)** ATP-linked respiration determined from the Mito Stress Test assay in AOA2 patient and carrier cells (n=5). **(E)** Representative graph of the Seahorse Glycolytic Stress Test assay of AOA2 patient and carrier cells measured on the y-axis as extracellular acidification rate (ECAR) and x-axis as time in minutes. Note the drugs used at different time points: glucose (gluc), oligomycin (oligo), and 2-Deoxy-D-glucose (2-DG). **(F)** Quantitation of basal extracellular acidification rate (ECAR) of AOA2 and carrier cells (n=5). **(G)** Quantitation of glycolytic reserve in AOA2 patient and carrier cells (n=5). Two-tailed paired t-tests with a 95% CI were used. P values: * = p < 0.05; ** = p < 0.01.

We next evaluated AOA2 patient cells for morphological differences in mitochondria and found that both carrier and patient cells displayed fused/filamentous mitochondria (Fig. S6, A and B). The mean fluorescent intensity (MFI) of TOM20 did not differ significantly between carrier and patient cells (Fig. S6, C and D). These results suggest that AOA2 patient cells may have normal mitochondrial morphology. However, abnormal mitochondrial morphology may be a phenotype of haploinsufficiency, as carrier cells harbor one inactive *SETX* allele. The frequencies of MN and lamin A/C blebbing did not differ significantly between patients and carriers and were very low (Fig. S6, D and E). These results suggest that the nuclear and genomic instability phenotypes do not extend to patient fibroblasts.

## Discussion

Our data show that *Setx^-/-^* MEFs exhibit elevated genomic and nuclear instability, potentially due to increased DNA damage (Fig. 1). This nuclear damage may promote a cGAS-STING-dependent type I IFN response (Fig. 2), though high ROS or mitochondrial damage may also contribute (Fig. 3). We also observe two novel metabolic phenotypes for *Setx^-/-^* cells: reduced mitochondrial membrane potential and increased ECAR rate. At the same time, mitochondria appear hyperfused and filamentous (Fig. 4, Fig. 5). ROS levels are also increased in *Setx^-/-^* cells (Fig. S2 A); this may be related to mitochondrial dysfunction, type I IFN induction, or both. Mitochondrial membrane potential is restored upon knockout of cGAS or *STING* in *Setx*^-/-^ MEFs, suggesting that innate immune activation via cGAS-STING drives mitochondrial dysfunction (Fig. 5). Both primary mouse brain tissue and AOA2 patient cells exhibit increased ISG expression. AOA2 patient cells also showed decreased mitochondrial function (Fig. 6, Fig. 7). Taken together, our data suggest a model in which cGAS and STING activate IRF3 to trigger an innate immune response, potentially due to increased cytosolic DNA from DNA damage-induced genomic and nuclear instability. This innate immune activation induces mitochondrial dysregulation (Fig. 8).

**Figure 8:**
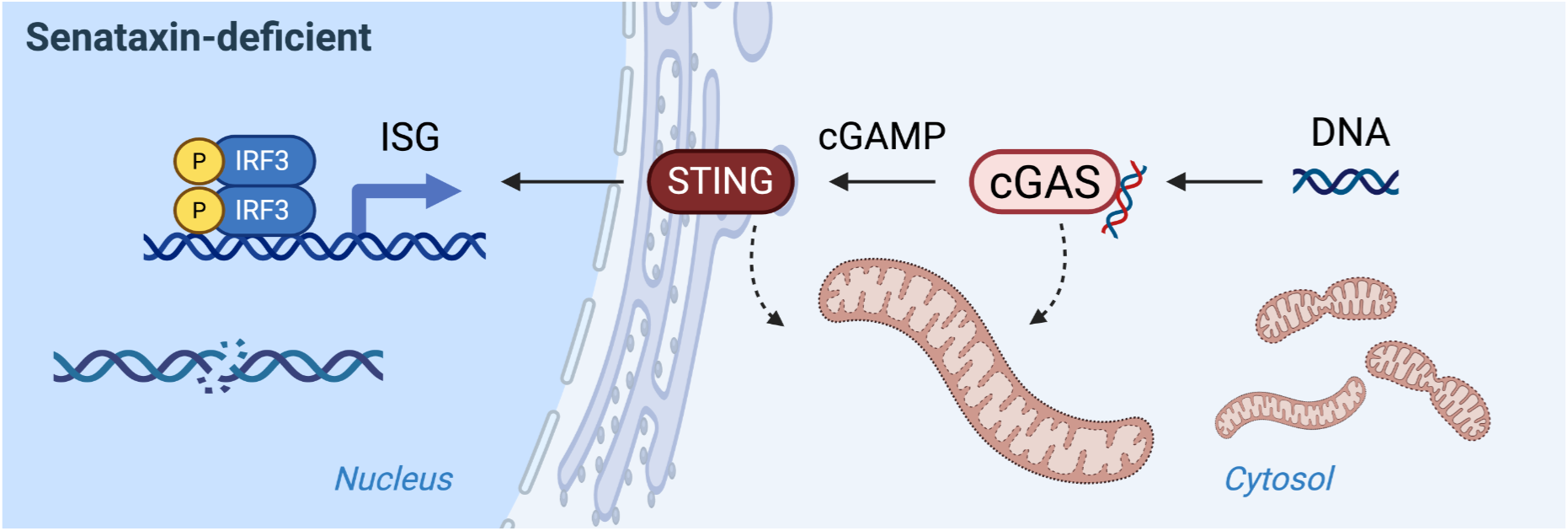
Working model of the cGAS-STING-IRF3 axis inducing ISG expression and mitochondrial depolarization in Senataxin-deficient cells. Here we report that cGAS-STING drives type I ISG expression and promotes mitochondrial dysfunction, likely inducing inflammation. We hypothesize that cytosolic DNA accumulates due to nuclear or mitochondrial instability, leading to cGAS-STING activation that either directly or indirectly causes mitochondrial depolarization. We speculate this results in further stress signaling via mtDNA release and induction of inflammation.

Innate immune activation and inflammation are common phenotypes in neurodegeneration and ataxias. An increase in cGAS expression has been observed in both patients and mouse models of Alzheimer’s disease (He et al., 2025). Mice lacking *C9orf72* expression, a known driver of amyotrophic lateral sclerosis (ALS) and frontotemporal dementia (FTD), also exhibit increased type I IFN responses and increased STING expression and activation (McCauley et al., 2020). In fragile X-associated tremor ataxia (FXTAS), there is a significant increase in inflammatory cytokines in patients’ brains, including IL-12 and TNF-α (Dufour et al., 2021). FXTAS patient fibroblasts also exhibit a significant increase in the type I IFN immune response (Napoli et al., 2021). Studies of FA using frataxin-deficient mice have also demonstrated increased inflammation, including hyperactivation of microglial cells (Shen et al., 2016). In A-T patients, proinflammatory cytokines such as IL-8 are also elevated (McGrath-Morrow et al., 2010). Further, A-T patient fibroblasts have increased innate immune activation induced by cGAS and STING, and inhibition of cGAS and STING reduces this activation (Aguado et al., 2021).

The Wolvetang group has shown that in A-T patient iPSC-derived brain organoid models, increased innate immune activation, mitochondrial dysregulation, and senescent phenotypes are observed (Aguado et al., 2021; Leeson et al., 2024). In one study, they showed that inhibition of cGAS and STING reduced inflammation, senescence, and neurodegenerative phenotypes in A-T brain organoid models (Aguado et al., 2021). They did not explore how cGAS and STING inhibition could influence the mitochondrial dysregulation they observe in A-T patient-derived organoids, but suggested that cGAS and STING inhibition may also alleviate mitochondrial phenotypes (Leeson et al., 2024). Similarly, we propose that inhibition of cGAS and/or STING is a promising therapeutic approach for AOA2 and may alleviate not only inflammatory phenotypes but also mitochondrial defects.

Many ataxias also show signs of oxidative stress, including A-T (Leeson et al., 2024) and FA (Lupoli et al., 2018). Due to its prevalence in ataxia and its ability to cause DNA damage, many speculate that increased ROS may drive ataxic initiation and/or progression (Ajayi et al., 2012; D’Souza et al., 2013; Lupoli et al., 2018; Palau & Espinós, 2006). Indeed, antioxidant compounds can reduce motor and cardiac phenotypes of FA in a mouse model and human patient stem cells (Villa 2021). For FA patients, the antioxidant drug Skyclarys (omaveloxolone) is FDA-approved. Both yeast lacking the Senataxin homolog, Sen1, and human HeLa cells with decreased *SETX* expression showed elevated ROS levels (Renaudin et al., 2021; Sariki et al., 2016); however, the source of the elevated ROS remains unknown. We also demonstrated increased ROS production in *Setx^-/-^* MEFs. Both innate immune activation (Hayman et al., 2021; Kumova & Carey, 2020) and mitochondrial dysfunction (Guo et al., 2013) can induce excessive ROS production. Dissecting the roles of innate immune activation and metabolism in ROS production, and how ROS, in turn, influence these pathways, will help determine the underlying phenotypes that may drive cerebellar degeneration in AOA2.

Mitochondrial dysfunction is a common hallmark of neurodegeneration (Klemmensen et al., 2023) and has been documented in FA (Abeti et al., 2016), A-T (Quek et al., 2017; Valentin-Vega et al., 2012), FXTAS (Ross-Inta et al., 2010), and AOA1 (Zheng et al., 2019). Here, we show that AOA2 patient-derived fibroblasts also exhibit mitochondrial dysfunction, as evidenced by decreased mitochondrial membrane potential and basal OCR. More comprehensive approaches, such as metabolomics, will be useful to determine the specific metabolites and metabolic pathways affected by the absence of Senataxin. In MEFs, we observe morphological differences in mitochondria, and OPA1 may be influencing them. However, many other factors can influence mitochondrial morphology, including deacetylation and SUMOylation of OPA1 (Yao et al., 2024), ubiquitination of MFN1 (Escobar-Henriques, 2014), and posttranslational modifications of FIS1 (Qin & Xi, 2022). Further work dissecting changes in mitochondrial remodelers will explain why the loss of Senataxin leads to morphological differences in MEFs and may help define the underlying cause of mitochondrial dysregulation in AOA2.

A limitation of our work is the use of patient-derived fibroblasts from one patient and a carrier. However, a study using weighted gene co-expression network analysis demonstrated that transcriptional profiles from blood samples of 32 patients from 20 families, the largest transcriptional study ever performed for this disease, exhibit a distinct AOA2 profile (Ngo et al., 2025). Further, transcriptional profiles from patient blood samples have been shown to be preserved in both patient fibroblasts and *Setx^-/-^*mice (Fogel, Lee, et al., 2014; Ngo et al., 2025). This work suggests that molecular phenotypes across samples are likely to be similar. Therefore, the AOA2 patient cells used in this study are representative of AOA2, but our findings should be further validated in additional AOA2 patient cell lines to determine the penetrance and range of mitochondrial dysfunction in affected individuals. Both AOA2 patient and carrier cells showed fused mitochondria; therefore, it is unclear if this phenotype is specific to MEFs or may also occur from haploinsufficiency. Future work using human fibroblasts with two functional copies of *Senataxin* would be highly useful for further examination of mitochondrial phenotypes. Further, it will be critical to validate these findings in additional physiological models, including experiments in mice, organoid models, or iPSCs differentiated into cerebellar neuronal populations, such as Purkinje cells, the dominant cell type in the cerebellum.

In summary, we demonstrate that in Senataxin-deficient cells, cGAS and STING promote mitochondrial membrane potential defects, as their deletion can reverse this phenotype. We further show that innate immune activation is present in *Setx^-/-^* mice and AOA2 patient fibroblasts, and that patient fibroblasts also show decreased mitochondrial membrane potential and OCR.

## Acknowledgements

We would like to thank Professors Frederic Chédin, Gino Cortopassi, Flora Tassone, and Natalie Niemi, as well as the members of the Chédin and Cortopassi labs for helpful conversations. Special thanks to Professor Brent Fogel for the rare carrier and AOA2 patient materials, and scientific and clinical insights into AOA2. We would also like to thank Drs. Priya Shah, Satoshi Namekawa, Neil Hunter, and Chang-il Hwang for sharing equipment and resources. Thank you to Jack McTiernan for assistance in figure design and layout. This project was supported by the National Ataxia Foundation Graduate Research Fellowship to JLAF, UC Davis Bridge funding, W.M. Keck Foundation (CRM0133094), and NIGMS R01 (GM134537) to Dr. J.H.B. This work was also supported by the University of California Davis Flow Cytometry Shared Resource Laboratory with funding from the NCI P30 CA093373 (Comprehensive Cancer Center), and technical assistance from Bridget McLaughlin, Jonathan Van Dyke, and Ashley Karajeh. Additionally, the UC Davis Drug Discovery Unit run by Dr. Alexey Tomilov in Dr. Gino Cortopassi’s lab provide guidance, support, and facilitated use of the Agilent Seahorse XFe96.

## Notes

### Competing Interest Statement

The authors have declared no competing interest.

