## Supplementary figures for "Senataxin loss induces cGAS–STING-mediated mitochondrial dysfunction"

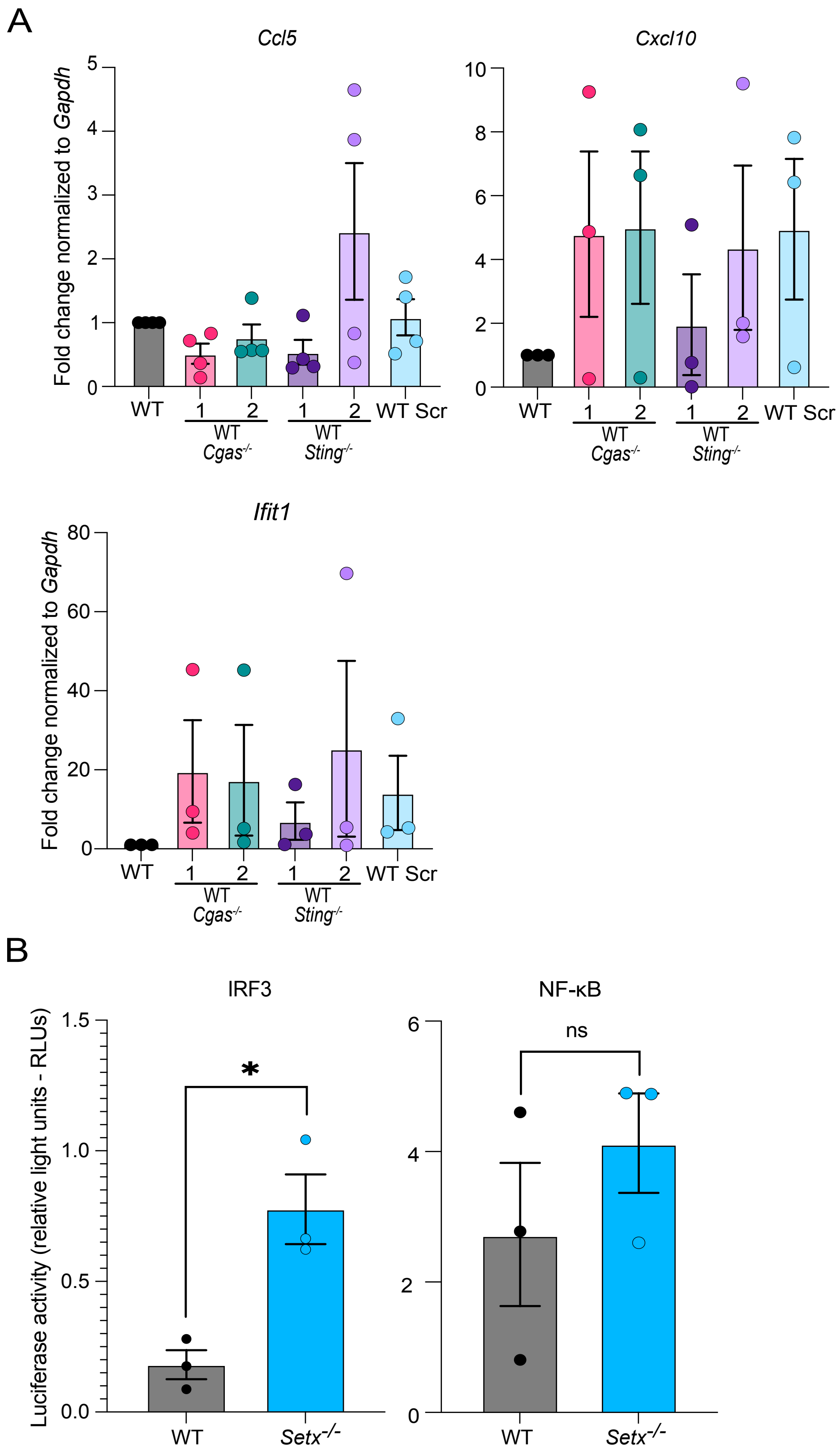

**Figure S1: *Cgas*$^{-/-}$ and *Sting*$^{-/-}$ do not significantly impact ISG expression, and Setx$^{-/-}$ MEF IRF3 activity is increased. (A)** ISG expression in WT, *Cgas*$^{-/-}$, and *Sting*$^{-/-}$ MEFs (n=3-4). **(B)** Dual luciferase reporter assay of IRF3- and NF-κB-dependent luciferase expression normalized to Renilla luciferase (n=3). One-way ANOVA (which graph) and 95% CI for two-tailed unpaired t-tests (which graphs). P values: * = $p < 0.05$.

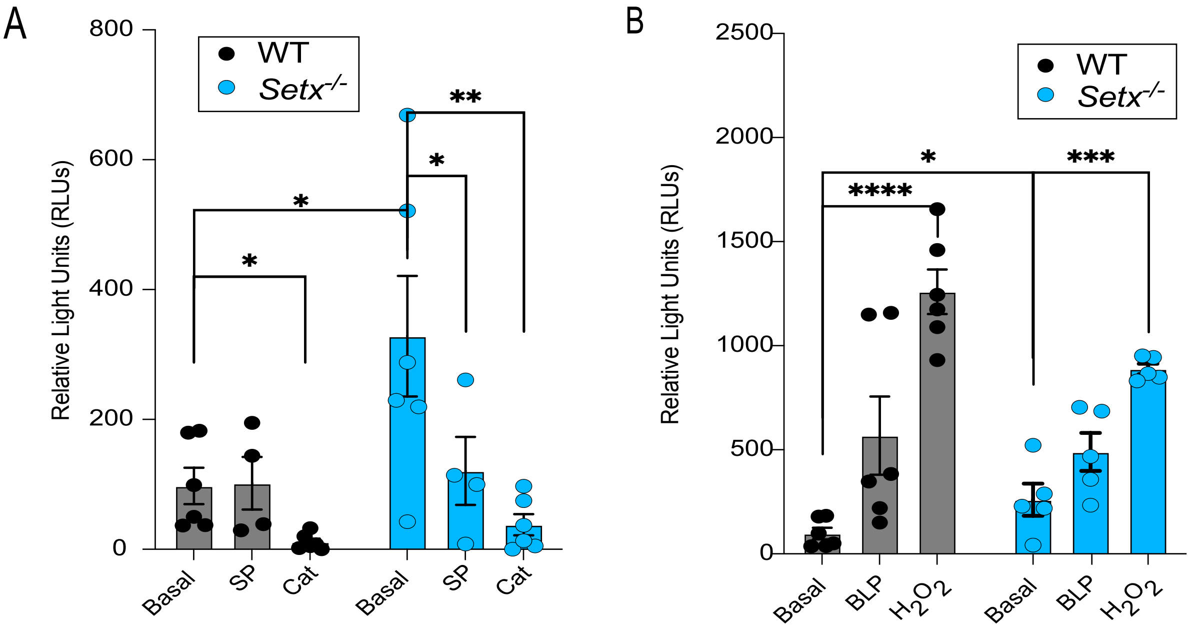


**Figure S2: ROS levels can be modulated with ROS scavengers and inducers. (A)** Quantitation of HRP-luciferase signal by 25 µM sodium pyruvate (SP) and 100,000 U/mL catalase (cat) in WT and *Setx*$^{-/-}$ MEFs (n=4-6). **(B)** Quantitation of HRP-luciferase signal in response to ROS inducers, 50 µM $H_2O_2$ and 1 µM beta-lapachone (BLP) in WT and *Setx*$^{-/-}$ MEFs (n=5-6). Two-tailed unpaired t-tests with 95% CI were used. P values: * = $p < 0.05$; ** = $p < 0.01$; *** = $p < 0.001$; **** = $p < 0.0001$.

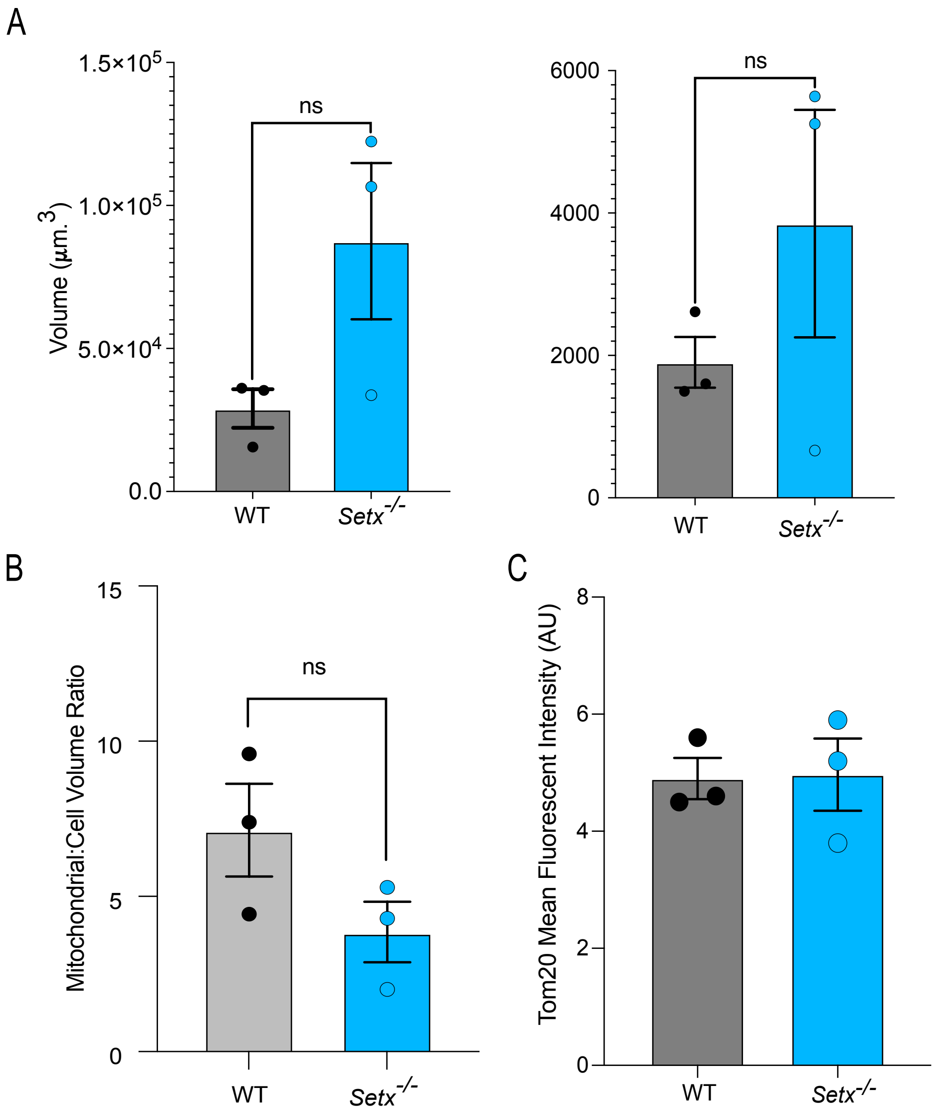


**Figure S3: *Setx*$^{-/-}$ MEFs' mitochondrial and cell volume proportions are similar to WT MEFs and have similar TOM20 MFI. (A)** Cell volume (left) and mitochondrial volume (right) in WT and *Setx*$^{-/-}$ MEFs (n=3). **(B)** The mitochondrial-to-cell ratio in WT and *Setx*$^{-/-}$ MEFs (n=3). **(C)** Quantitation of TOM20 MFI in WT and *Setx*$^{-/-}$ MEFs (n=3). Two-tailed unpaired t-tests with 95% CI were used. P values: ns = not significant.

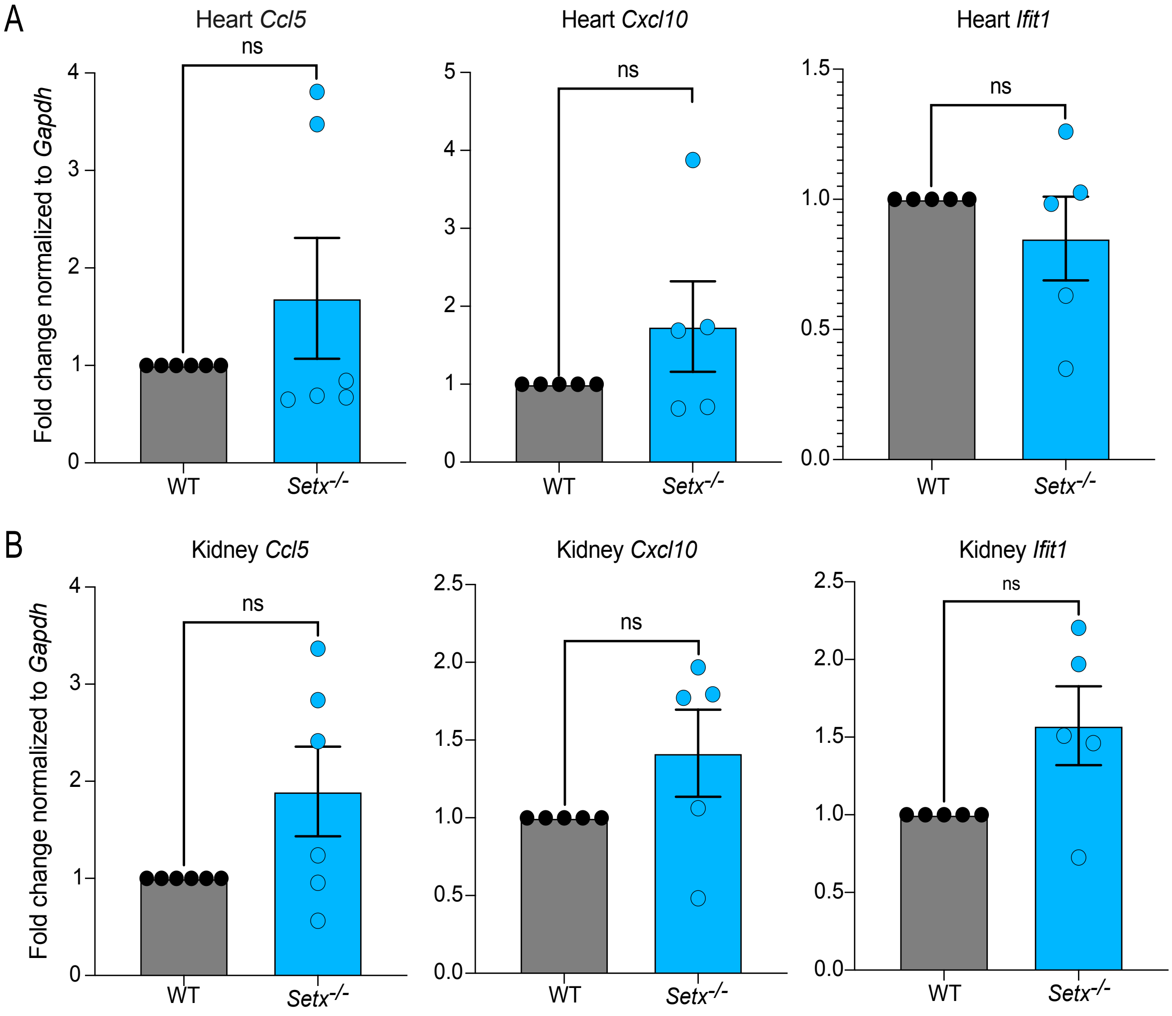


**Figure S4: ISG expression in hearts and kidneys from *Setx*$^{-/-}$ mice is similar to WT. (A)** ISG expression was measured by RT-qPCR in heart tissue from WT and *Setx*$^{-/-}$ mice (n=5-6). **(B)** ISG expression was measured by RT-qPCR in kidney tissue from WT and *Setx*$^{-/-}$ mice (n=5-6). Two-tailed unpaired t-tests with 95% CI were used. P values: ns = not significant.

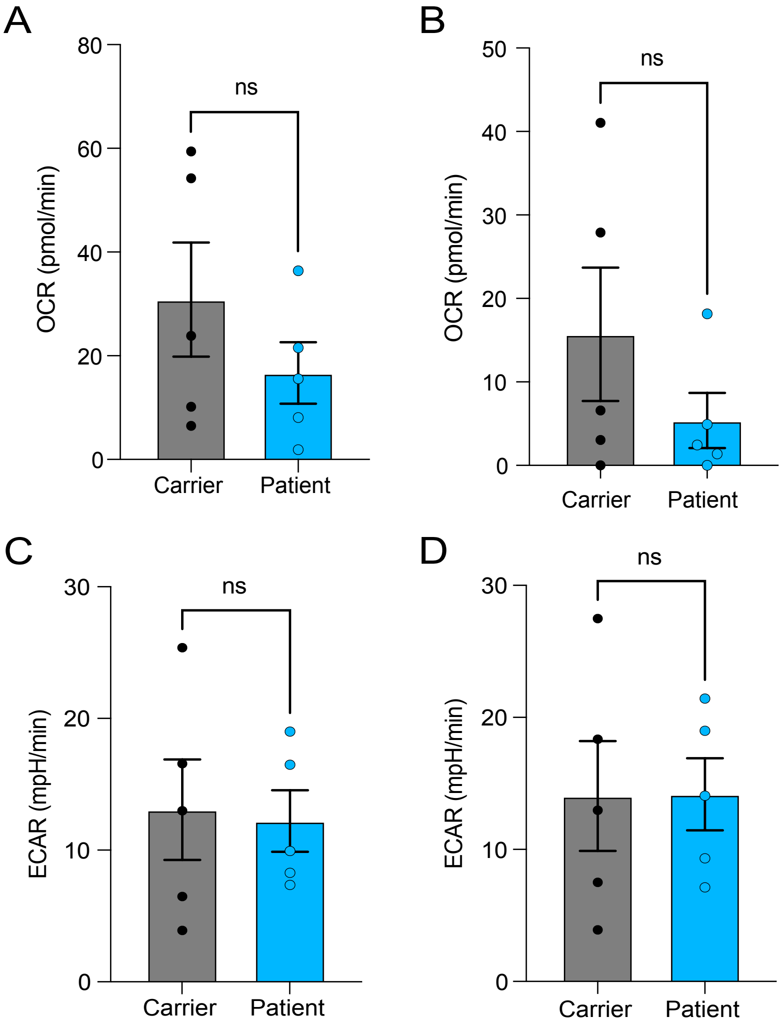


**Figure S5: Differences in respiratory and glycolytic measurements from Seahorse Mito Stress Test and Glycolytic Stress Test assays. (A)** Quantification of maximal respiratory capacity in patient and carrier cells using the Mito Stress test assay (n=5). **(B)** Quantitation of the spare respiratory capacity in patient and carrier cells using the Mito Stress test assay (n=5). **(C)** Quantitation of ECAR-attributed glycolysis in carrier and patient cells (n=5). **(D)** Glycolytic capacity as determined by ECAR changes in carrier and patient cells (n=5). Two-tailed paired t-tests with 95% CI were used. P values: ns = not significant.

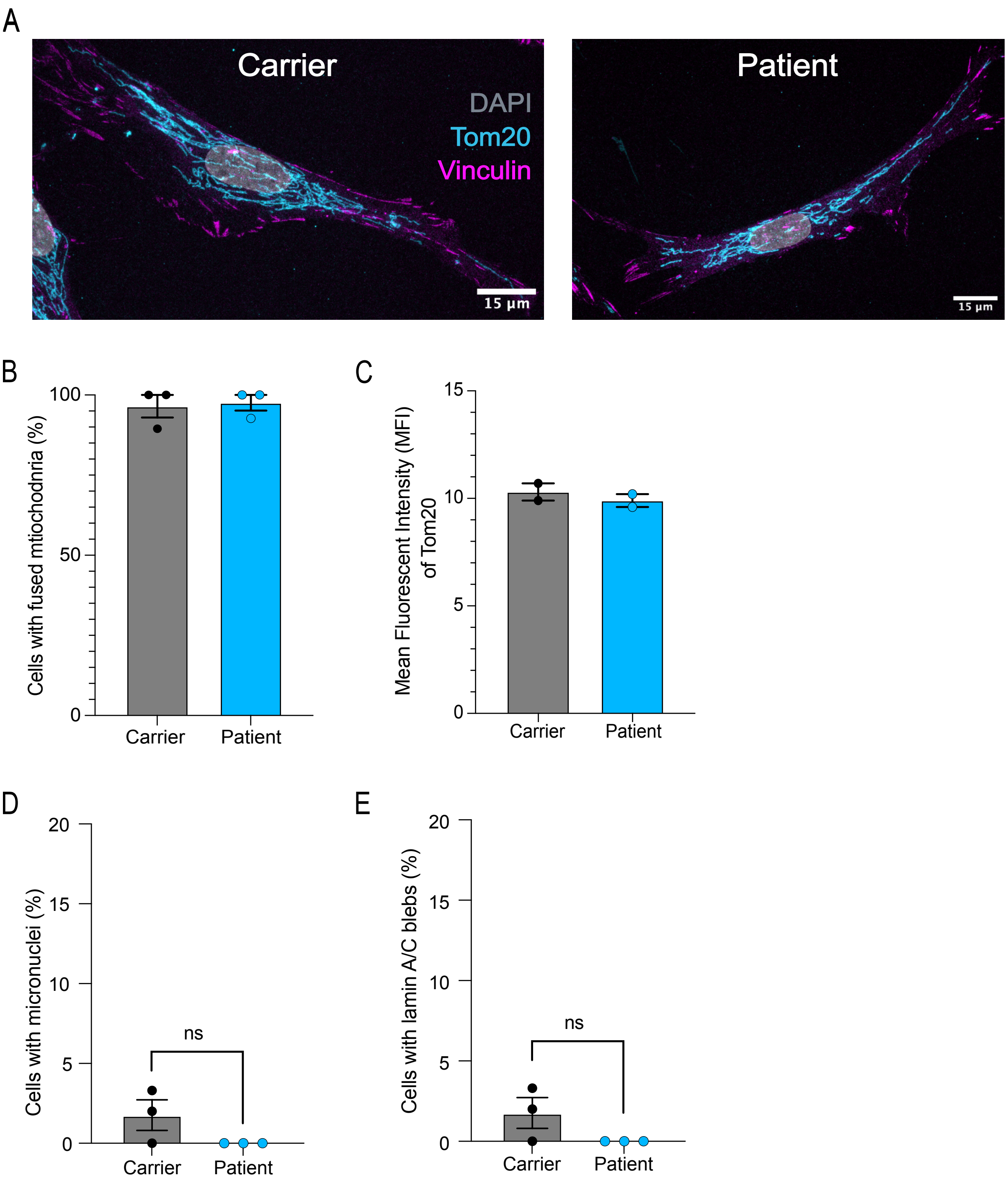


**Figure S6: AOA2 patient and carrier cells do not differ in morphology. (A)** Representative images of AOA2 carrier (left) and patient (right) cells labeled with DAPI (DNA, nucleus; grey), Tom20 (mitochondria; cyan), and Vinculin (cell membrane; magenta). **(B)** The percentage of cells with fused mitochondria in carrier and AOA2 patient cells (n=3). **(C)** The MFI of TOM20 signal in patient and carrier cells (n=2). **(D)** The percentage of cells with MN in carrier and patient cells (n=3). **(E)** The percentage of cells with lamin A/C blebs in carrier and patient cells (n=3). Two-tailed unpaired t-tests with a 95% CI were used. P values are ns (not significant).
